# Chromosome-scale assembly of the African yam bean genome

**DOI:** 10.1101/2023.10.31.564964

**Authors:** Bernice Waweru, Isaac Njaci, Edwin Murungi, Rajneesh Paliwal, Collins Mulli, Mary Maranga, Davies Kaimenyi, Beatus Lyimo, Helen Nigussie, Bwihangane Birindwa Ahadi, Ermias Assefa, Hassan Ishag, Oluwaseyi Olomitutu, Michael Abberton, Christopher Darby, Cristobal Uauy, Nasser Yao, Daniel Adewale, Peter Emmrich, Jean-Baka Domelevo Entfellner, Oluwaseyi Shorinola

## Abstract

Genomics-informed breeding of locally adapted, nutritious, albeit underutilised African crops can help mitigate food and nutrition insecurity challenges in Africa, particularly against the backdrop of climate change. However, utilisation of modern crop improvement tools including genomic selection and genome editing for many African indigenous crops is hampered by the scarcity of genetic and genomic resources. Here we report on the assembly of the genome of African yam bean (*Sphenostylis stenocarpa)*, a tuberous legume crop that is indigenous to Africa. By combining long and short read sequencing with Hi-C scaffolding, we produced a chromosome-scale assembly with an N50 of 69.5 Mbp and totalling 649 Mbp in length (77 - 81% of the estimated genome size based on flow cytometry). Using transcriptome evidence from Nanopore RNA-Seq and homology evidence from related crops, we annotated 31,614 putative protein coding genes. We further show how this resource improves anchoring of markers, genome-wide association analysis and candidate gene analyses in Africa yam bean. This genome assembly provides a valuable resource for genetic research in Africa yam bean.

## Background and Summary

African yam bean (*Sphenostylis stenocarpa* (Hochst. Ex. A. Rich) Harms) is an underutilised tuberous legume which produces edible protein-rich seeds and starch-rich tubers (Fig. 1). It is a tropical African crop^1^ that originated from Ethiopia from where its distribution extended to West and Central Africa^2^. African yam bean (hereafter referred to as AYB) is important for food and nutritional security in local communities in sub-saharan Africa. AYB is a rich source of dietary protein with up to 30% and 10% protein content in the seeds and tubers, respectively^3,4^. Its seeds and tubers are also low in fat and rich in carbohydrates, minerals and vitamins^3,4^. In addition, AYB exhibits high nitrogen-fixing ability^5^ and is drought tolerant. These attributes may have allowed it to thrive in marginal soils under low-input farming systems and intercropping, especially in Ghana and Nigeria^6,7^.

**Fig. 1:**
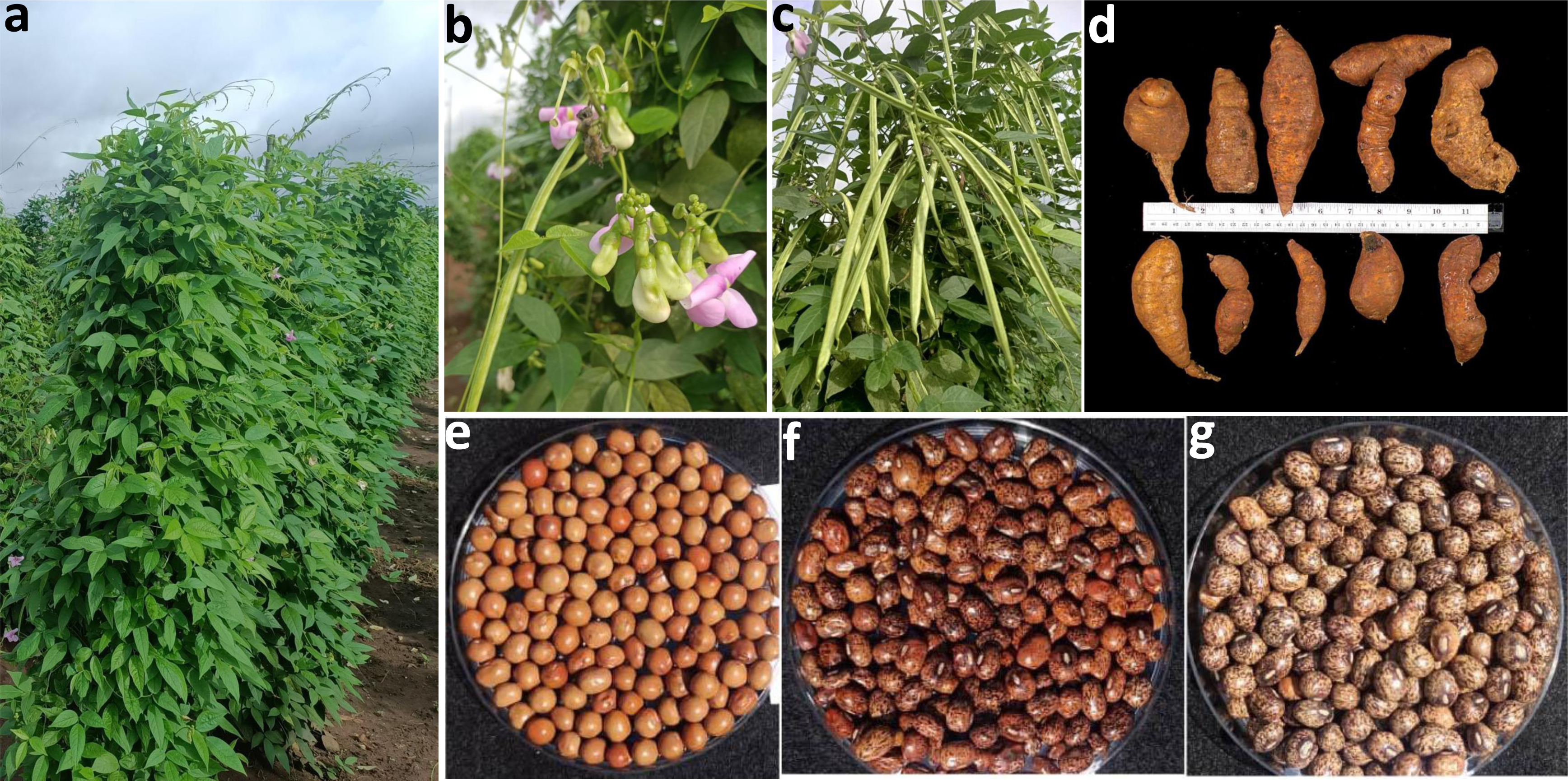
Africa yam bean - an African indigenous tuberous legume. Figure shows (a) full grown plants in the field (b) flowers (c) pods (d) root tubers of different shapes and sizes (e - g) different coloured seeds.

AYB, however, is largely underutilised due to the hardness of the seed coat, leading to long cooking time and the presence of anti-nutritional factors which reduce protein digestibility^3^. Also, the need for staking of plants has greatly hampered its cultivation on a commercial scale. Its production has been sustained indigenously through intercropping with major crops, especially yam - *Dioscorea spp*. To date, minimal genomic information is available to assist breeding efforts aimed at unlocking the full potential of AYB, thereby limiting its contribution to food and nutritional security in Africa.

Here, we present the first chromosome-scale assembly of the AYB genome using Illumina short-read and Oxford Nanopore long-read sequencing platforms (Fig. 2). Using homology and transcript evidence, we performed a gene annotation of the AYB’s genome (Fig. 2). We further demonstrate the usefulness of this genome resource for genetics analyses and trait mapping in AYB.

**Fig. 2:**
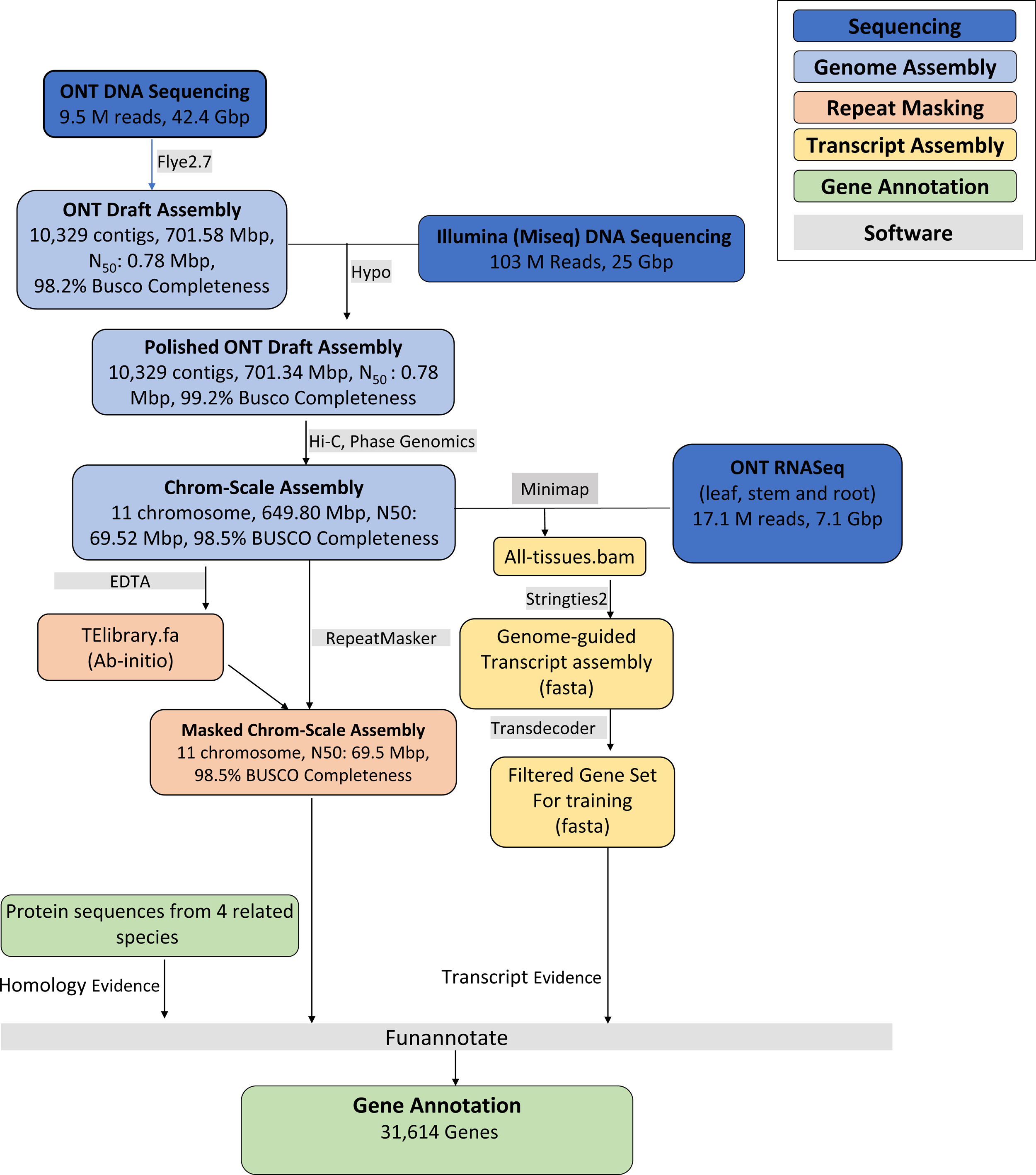
AYB genome sequencing and annotation workflow. Overview of the workflow used for the sequencing, assembly, masking and annotation of the African Yam Bean genome. The boxes are color-coded by the different stages involved in the workflow (top right box). Software used for each step are indicated in grey boxes..

## Methods

### Size estimation of AYB genome by flow cytometry

Fresh 10 mg leaf samples of AYB and soybean (*Glycine max*, used as standard) were immersed in 1 mL of ice-chilled Galbraith buffer (45 mM MgCl_2_, 30 mM sodium citrate, 20 mM 3-(N-morpholino) propanesulfonic acid, 0.1% w/v Triton X-100, pH 7) and sliced using a scalpel. The supernatant was filtered through one layer of Miracloth (pore size 22 - 25 µm). An aliquot of 600 µL of filtrate were mixed with propidium iodide to a concentration of 50 µM and RNAse A to 20 µg/mL and incubated for 1.5 h on ice. A FACSCantoll flow cytometer (Becton Dickinson) was used to measure nuclei, with flow rate adjusted to 20 - 50 events/s and results were analysed using FCSalyser (v. 0.9.18 alpha). The genome size of *Sphenostylis stenocarpa* was estimated following the method described by Dolezel et al (2007)^8^ by dividing the mean position of its fluorescence peak by the mean position of the corresponding soybean peak and multiplying by the estimated soybean genome size of 1.10 - 1.15 Gbp^9^ (Fig. 3, Supplementary Table 1). Based on this range we estimate the size of the AYB genome as 804 - 841 Mbp.

**Fig. 3.**
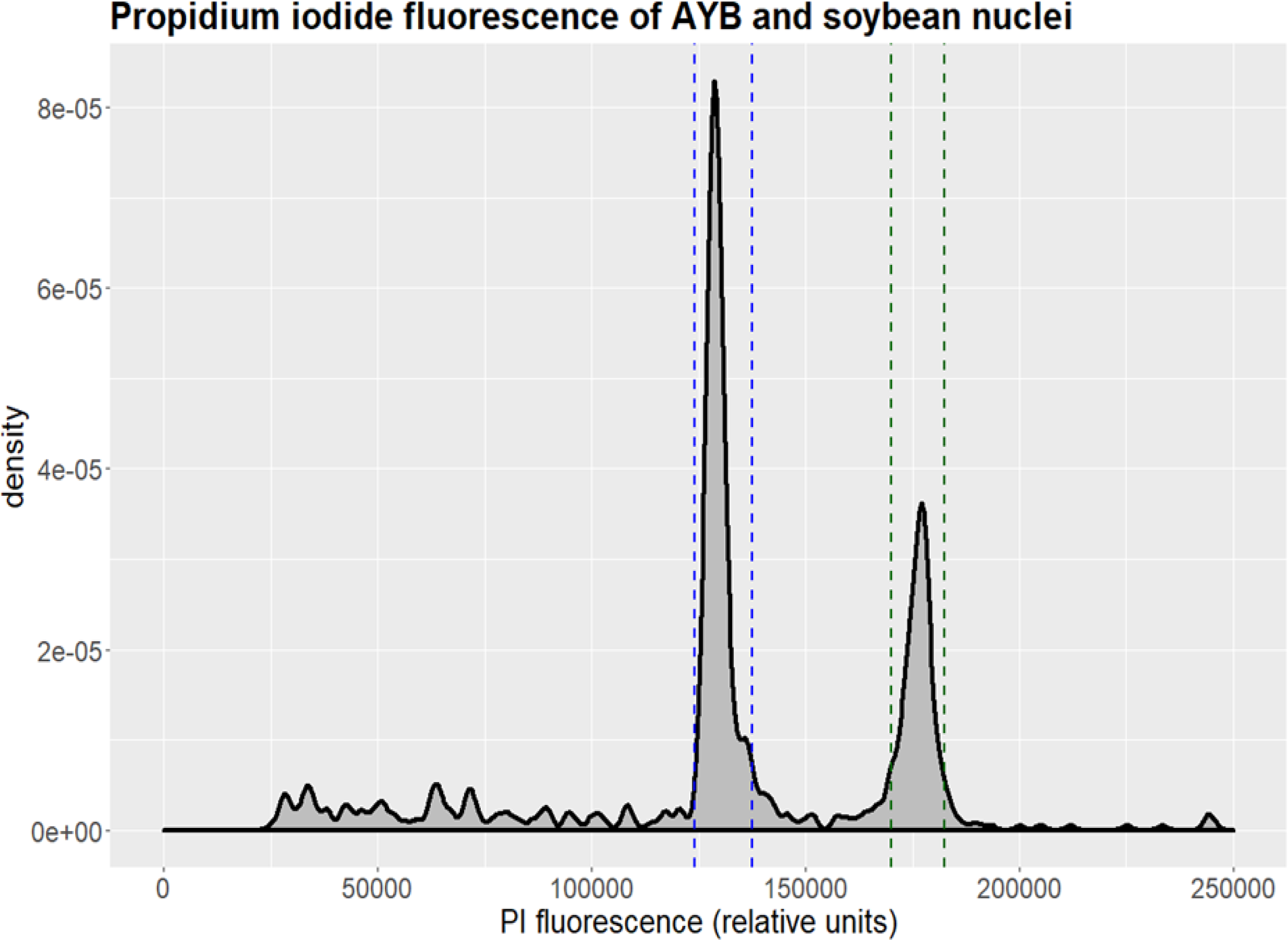
Genome size estimation of AYB. Density plot showing results of a representative flow cytometry run, excluding events caused by cell debris. Propidium iodide fluorescence amplitude (in relative units) is plotted against event density. The interval assumed to be AYB nuclei is delimited by blue dashed lines, the interval assumed to be soybean nuclei is delimited by green dashed lines. Three biological replicates were performed.

### Sample selection, library preparation and sequencing

#### Illumina DNA sequencing

Seeds of AYB accession TSs11^10^ were germinated in a Petri dish on filter paper moistened with tap water. The sprouted seedlings were transferred to soil and allowed to grow in the greenhouse facility at the International Livestock Research Institute (ILRI, Kenya) for a month. DNA was extracted from young leaves using a DNeasy Plant Mini Kit (Qiagen, Hilden, Germany) following the manufacturer’s protocol, recovering a total of 10 µg. DNA was quantified using a Qubit 2.0 Fluorometer and dsDNA BR Assay (Invitrogen, Paisley, United Kingdom) and integrity was confirmed by gel electrophoresis on a 0.8% agarose gel.

Three samples of 50 ng each of genomic DNA were sheared and processed using the Nextera DNA Library Prep Kit (Illumina, USA) according to the manufacturer’s instructions. Three runs of paired-end (2 x 150 bp) sequencing were performed on an Illumina MiSeq (Illumina) to generate 25 Gbp of raw data, representing ∼30x of the AYB genome.

#### Nanopore DNA sequencing

Two grams of young leaves of AYB accession TSs11 were harvested, frozen in liquid nitrogen and stored at −80 ^0^C. The leaves were then ground in liquid nitrogen using a pestle and mortar. High Molecular Weight (HMW) DNA was extracted from the ground sample with Carlson lysis buffer (100 mM Tris-HCl, pH 9.5, 2% CTAB, 1.4 M NaCl, 1% PEG 8000, 20 mM EDTA) followed by purification using the Qiagen Genomic-tip 100/G as described on the Oxford Nanopore Technologies (ONT, UK) HMW plant DNA extraction protocol. The ONT SQK-LSK109 ligation sequencing kit protocol was used to prepare sequencing libraries from the HMW DNA. This involved repairing and 3’ adenylation of 1 µg of HMW genomic DNA with the NEBNext FFPE DNA Repair Mix and the NEBNext® Ultra™ II End Repair/dA-Tailing Modules (New Englang Biolab, NEB). Sequencing adapters were then ligated using the NEBNext Quick Ligation Module (NEB). After library purification with AMPure XP beads (Beckman Coulter), sequencing was conducted at ILRI using R9.4.1 flow cells on an ONT MinION sequencer. High-accuracy base calling was done using Guppy basecaller^11^ (v4.1.1) generating 9.5 million reads totalling 42.4 Gbp of sequence that represents 50 - 53x of the estimated genome size (Fig. 2).

#### Nanopore RNA sequencing

Two grams of young and disease-free leaves, stem and root tissues of AYB accession TSs11 were harvested and ground with mortar and pestle in liquid nitrogen. HMW RNA was extracted with the following extraction buffer [100 mM Tris–HCl (pH 8.0), 25 mM EDTA, 2 M NaCl, 2% CTAB (w/v), 2% PVP (w/v) and 2% β-mercaptoethanol (v/v)], followed by removal of residual DNA using DNASE I (RNase-free) kit (Thermo Fisher Scientific). The library was prepared following the Oxford Nanopore SQK-PCS109 PCR-cDNA sequencing kit. A total of 50 ng total RNA was transcribed using Maxima H Minus Reverse Transcriptase (Thermo Fisher Scientific). Full length transcripts were selected by PCR amplification using the LongAmp Taq Master Mix and the product was purified with AMPure XP beads (Beckman Coulter). An aliquot of 1 µL of Rapid Adapter was added to the amplified cDNA library. The libraries were sequenced at ILRI using R9.4.1 flowcells on the ONT MinION sequencers. Real-time data acquisition and high accuracy base-calling were conducted using the MinKNOW software with the Guppy basecaller generating 7.1 Gbp of sequence from 17.1 million reads (Fig. 2).

### De novo assembly

Genome assembly was done primarily with the ONT long reads generated above. Briefly, the reads were assembled using Flye *de novo* long read assembler (v2.9)^12^ with default parameters generating 10,329 contigs with total assembly length of 701.6 Mbp (Fig. 2). The draft assembly was further polished for error correction with Illumina short reads generated above from the same AYB accession TSs11 using HyPo hybrid polisher^13^ (v1.0.3) with parameters -s 700m -c 30 -p 96 and -t 64. The polished draft assembly had an N50 of 781,337 bp, and a total assembly of 701.3 Mbp (Table 1). The draft assembly was further scaffolded using Chromatin conformation capture data as described below.

**Table 1:** AYB assembly statistics before and after scaffolding.

| Assembly Metric | Polished Assembly | Scaffolded (Hi-C) Assembly |
| --- | --- | --- |
| **Number of contigs/scaffolds (> 500 bp)** | 10,329 | 11 |
| **Total assembly length (bp)** | 701,349,621 | 649,801,261 |
| **N50 (bp)** | 781,088 | 69,519,929 |
| **L50(bp)** | 194 | 4 |
| **N75 (bp)** | 292,358 | 47,252,539 |
| **L75(bp)** | 567 | 7 |
| **Longest contig/scaffold (bp)** | 9,386,731 | 107,191,003 |
| **Mapping back rate of ONT reads** |  | 97.82% |

### Hi-C scaffolding

Chromatin conformation capture (Hi-C) scaffolding was performed by Phase Genomics (Seattle, USA) using the Proximo Hi-C 2.0 Kit. For this, fresh leaves from young AYB accession TSs11 plants were frozen in liquid nitrogen, ground to powder and cross-linked using formaldehyde solution before being sent to Phase Genomics for library preparation following the manufacturer’s protocol. Sequencing of the Hi-C library was performed at Phase Genomics using Illumina HiSeq platform generating a total of 275,166,448 paired-end reads. Reads were aligned against the polished assembly using BWA-MEM^14^ with options -5SP and -t 8 specified and the other parameters set to the defaults. PCR duplicates were then mapped using SAMBLASTER^15^, which were later excluded from the analysis. Non-primary and secondary alignments were flagged and filtered with Samtools^16^ using the -F 2304 filtering flag. Putative misjoined contigs were broken using Juicebox^17^ based on the Hi-C alignments, and the same alignment procedure was repeated from the beginning on the corrected assembly. Phase Genomics Proximo Hi-C genome scaffolding platform was used to create chromosome-scale scaffolds from the corrected assembly following the method similar to that described by Bickhart et al (2017)^18^. Ordering of the scaffolds into pseudomolecules was done by LACHESIS^19^.

The scaffolded assembly contains 11 pseudomolecules (representing the 11 AYB chromosomes) with 649.8 Mbp of sequence, and N50 of 69.5 Mbp (Table 1). There were also 8,422 short contigs with total and average length of 51.8 Mbp and 6.1 Kbp, respectively, that were not anchored into chromosomes. Summary statistics and evaluation of the completeness of the chromosome-scale genome assembly was evaluated using QUality ASsessment Tool (QUAST)^20^ (ver 5.0.2) and Benchmarking Universal Single-Copy Orthologs (BUSCO)^21^ (v5.2.2), respectively (see Technical Validation section).

### Synteny with closely related species genomes

We examined the syntenic relationship between the HiC-scaffolded genome of AYB and those of closely related species including common bean (*Phaseolus vulgaris)*^22^ and lablab (*Lablab purpureus)*^23^. For this, long-read based genome assemblies and annotation dataset for *P. vulgaris* and *L. purpureus* were obtained from Ensembl plant^24^ and e!DAL^25^ respectively. The blastp option from BLAST v2.7.1 was used to compare the AYB protein to *P. vulgaris* and *L. purpureus* with parameters: - max_target_seqs 1 -evalue 1e-10 -qcov_hsp_perc 70. MCscanX^26^ algorithm was subsequently used to identify collinear blocks between the AYB-phaseolus and AYB-lablab genome pairs with parameters: -s 20 and -m 10. Visualisation of synteny linkages was made by R^27^ (v3.3.1) and circos^28^ (v0.69-4). Six of AYB chromosomes show direct one-to-one syntenic relationships with lablab and common bean chromosomes, while the other five AYB chromosomes show syntenic relationship across two or more chromosomes in either lablab or common bean. Based on these syntenic relationships, we assigned chromosome names to the Hi-C-scaffolded AYB pseudomolecules.

### Repeat annotation

The Extensive de novo TE Annotator^29^ (EDTA v1.9.7) pipeline was used to annotate the transposable elements (TE) in the AYB genome. The pipeline incorporates different tools to annotate predominant TE classes found in plant genomes using structure and homology-based detection methods. The tools include LTRharvest^30^, LTR_FINDER^31^, LTR_retriever^32^, TIR-Learner^33^, HelitronScanner^34^, RepeatModeler2^35^ and RepeatMasker^36^. The outputs of each tool are combined and filtered into a comprehensive non-redundant TE library. The inbuilt genome annotation function in EDTA was then used to produce a final non-overlapping repeat annotation for the AYB genome. Data visualisation and summary were carried out in R^37^ using the Tidyverse suite^38^. In total 624,517 TEs and 78,100 unclassified repeats accounting for 74.08% of the total assembly were identified (Table 2, Fig. 5 and Fig. 6).

**Table 2:**
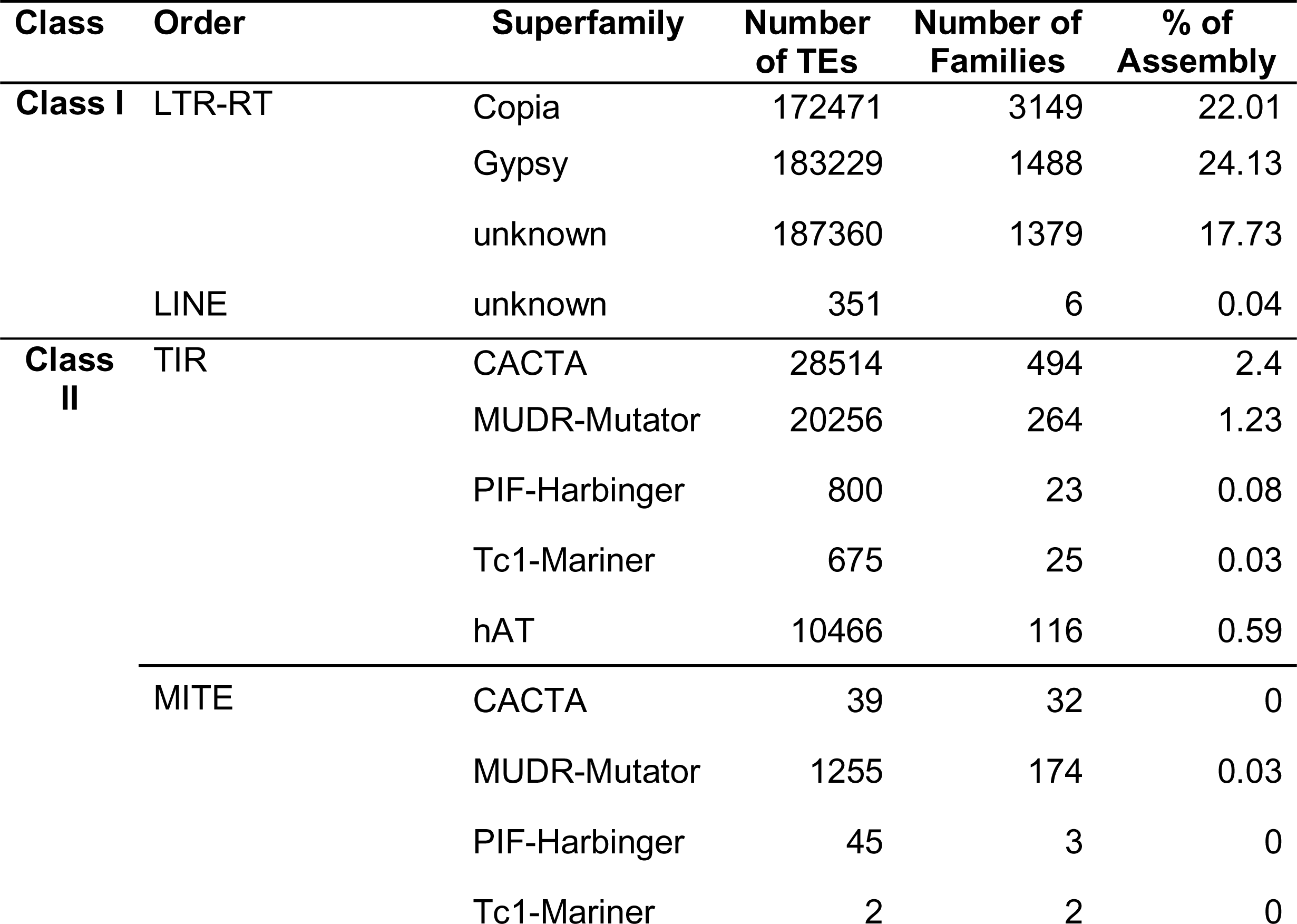
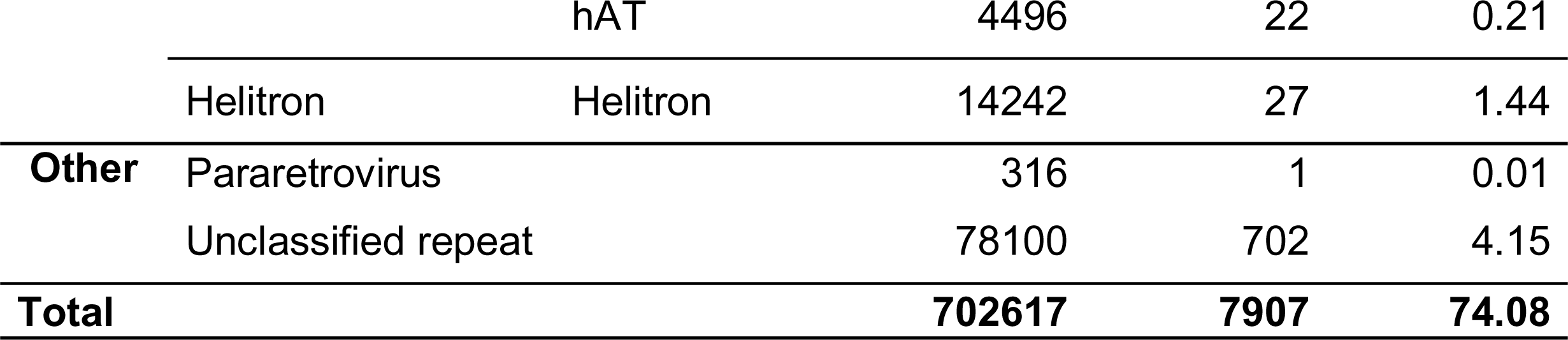
The number of TEs, TE families and the proportion of occupied assembly length by different classes of repeats identified and annotated in AYB.

**Fig. 4:**
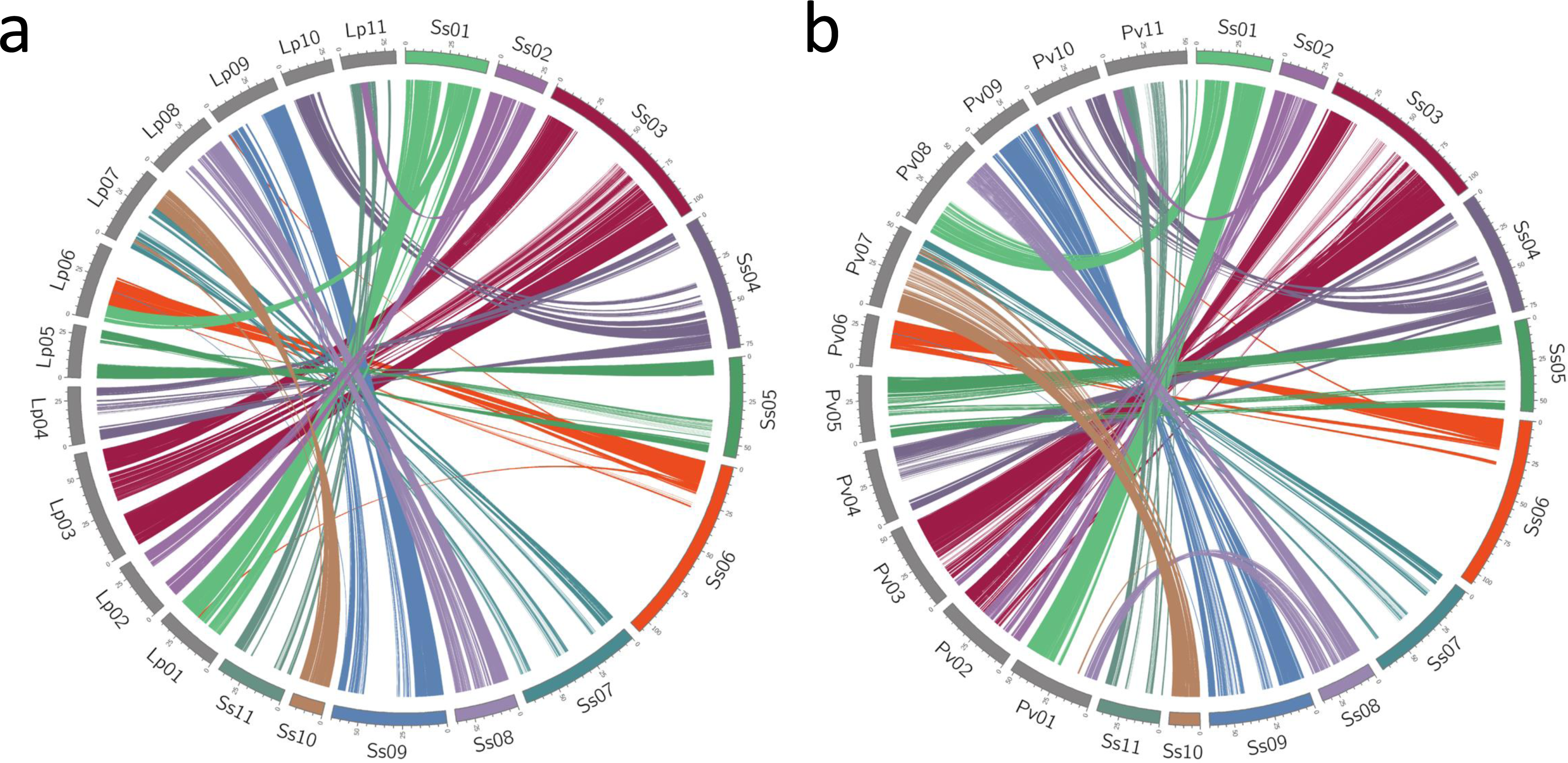
AYB synteny to related legumes. Syntenic relationships between AYB chromosomes to the genomes of (a) lablab (*Lablab purpureus*) and (b) common bean (*Phaseolus vulgaris*). Where possible, AYB chromosome were renamed to reflect these syntenic relationships.

**Fig. 5:**
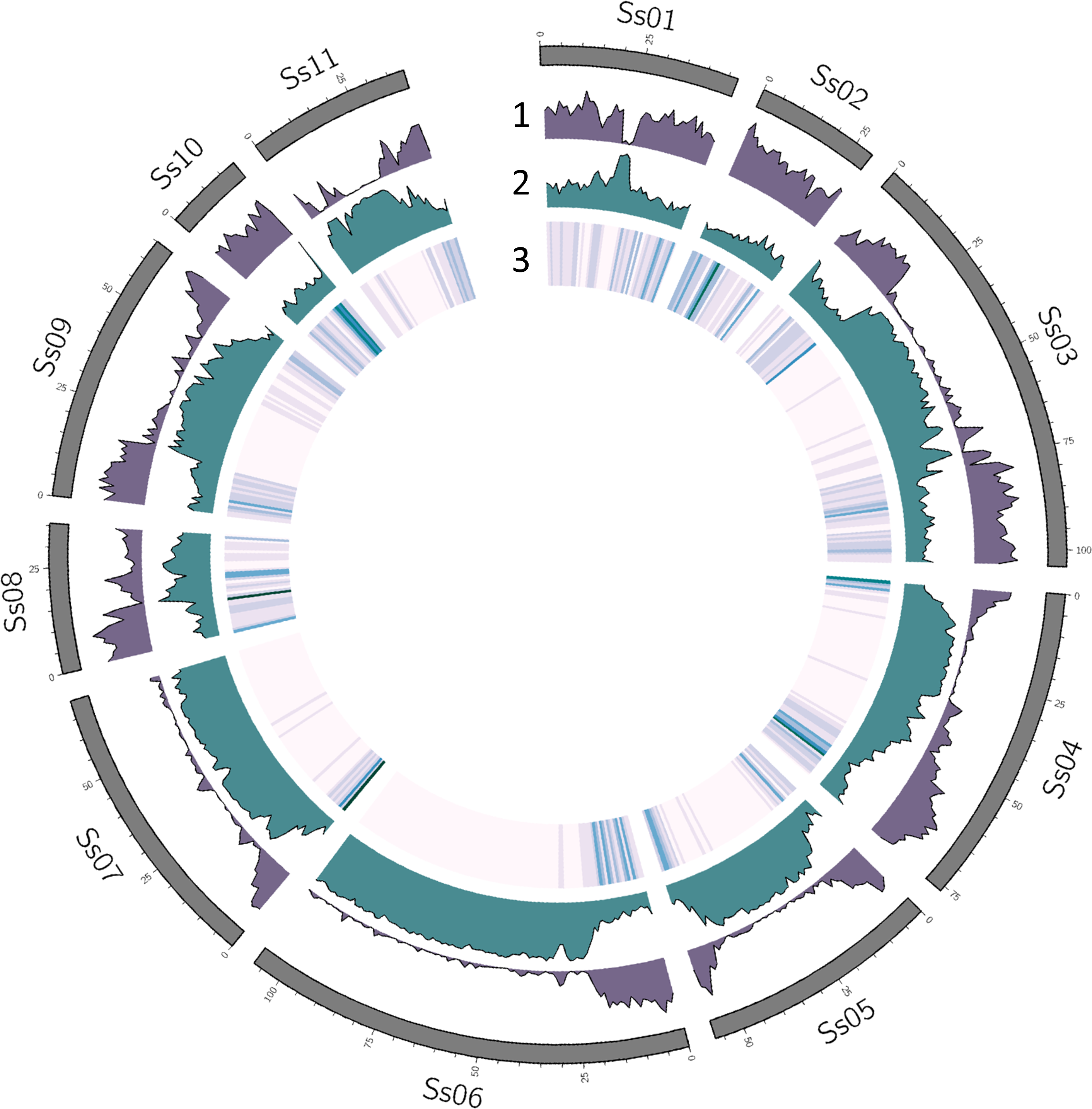
Gene, repeat and marker distribution in the AYB genome. The outer to the inner track show 1) gene density, 2) repeat density, 3) heatmap of marker distribution.

**Fig. 6:**
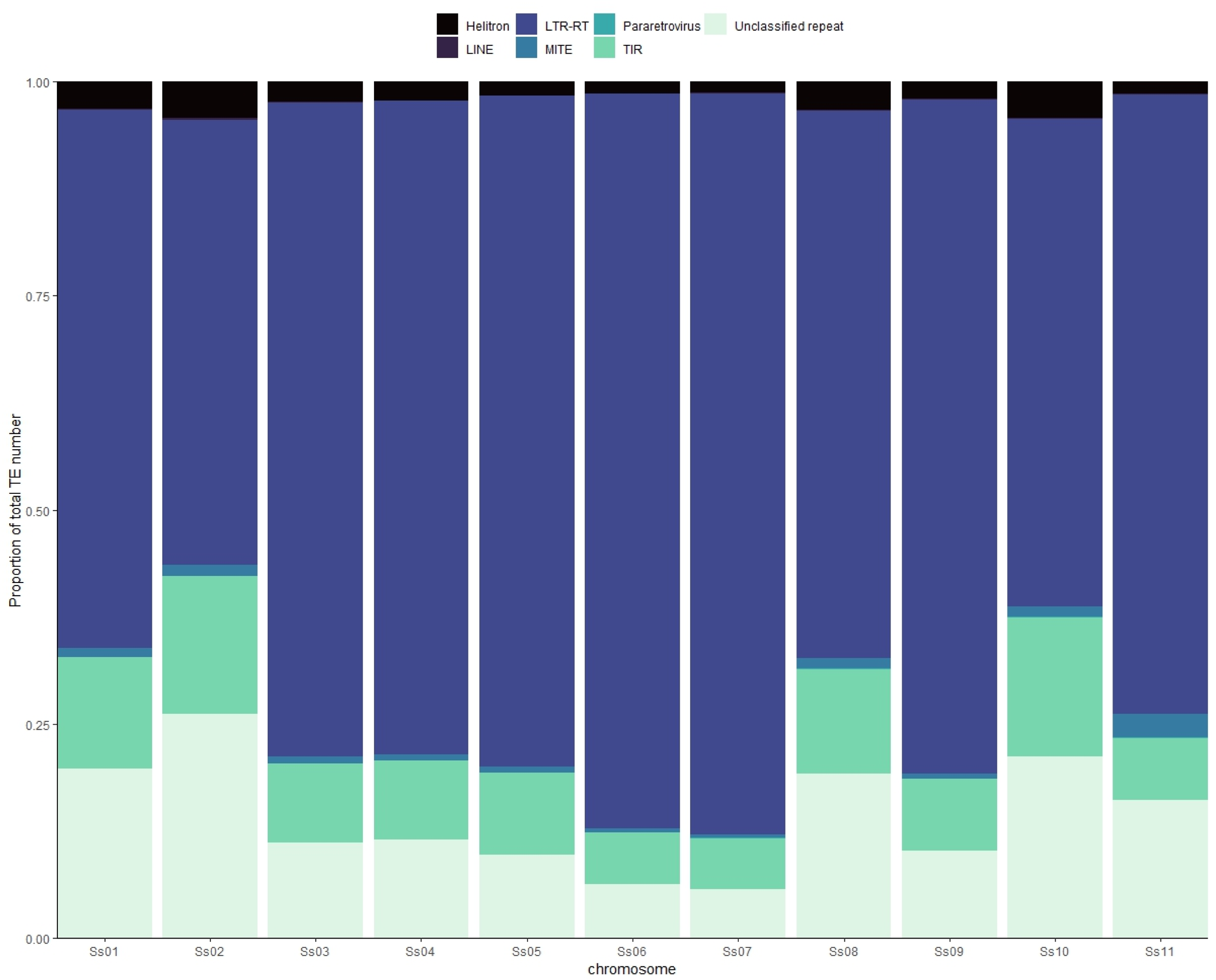
Distribution of transposable elements across the AYB genome. Chromosomal repeats content in the AYB genome showing proportional abundance of identified transposable element Orders on each chromosome.

### Gene prediction and functional annotation of genome

We combined transcript and homology evidence to annotate the gene content of the AYB genome. The transcript evidence was generated from 17,117,377 ONT long-read RNA reads totalling 7.1 Gbp of sequencing data used for *de novo* assembly of 60,249 transcripts. Briefly, Minimap2^39^ (v2.22) was used to index the AYB genome assembly and the RNA reads were mapped to the indexed assembly. Samtools^16^ (v1.9) was used to sort mapped reads by coordinates that were used to assemble transcripts with Stringtie2^40^ (v2.0.1). Transdecoder^41^ (v2.0.1) was then used to identify candidate CDS regions and select transcripts with a minimum protein length of 100 amino acids.

We combined the *de novo* transcripts with protein homology evidence from four well-annotated plant genomes (*Arabidopsis thaliana* TAIR10, *Phaseolus vulgaris* v1.0, *Glycine max* v2.1, *Vigna_angularis* v1.1) together with a soft-masked (for repeats) AYB genome as inputs into Funannotate^42^ (v1.8.11) to identify protein coding genes. Funannotate *‘predict’* uses *ab initio* gene predictors Augustus^43^, PASA^44^, SNAP^45^ and GlimmerHMM^46^ together with protein sequences as evidence to predict genes. Gene predictions from all four *ab initio* predictors are passed to EVidenceModeler^47^ with various weights for integration. This resulted in 30,840 coding gene models totalling 31,614 transcripts with a median exon length of 231 bp and a median of three exons per transcript. Additionally, we detected 774 non-overlapping tRNA-encoding genes using tRNAscan-SE^48^ for tRNA prediction. The gene and transposable element distribution across the genome are inversely correlated (Fig. 5).

Protein domains were annotated using InterProScan-5.25-64.0^49^ based on InterPro protein databases, including TIGRFAM, SUPERFAMILY, PANTHER, Pfam, PRINTS and ProDom. We also used eggNOG-mapper^50^ (v2.1.7) to annotate predicted gene models. Funannotate *‘annotate’* uses results of InterProScan and eggNOG-mapper to annotate putative functions of protein sequences using PFAM^51^, UniProtKB^52^ and Gene Ontologies^53^ databases. In total, functional descriptions were assigned to 25,241 (81.85%) of the genes.

### Gene family analysis

To delineate gene families, AYB was compared to other legumes *L. purpureus*, *P. vulgaris, Vigna unguiculata,* and *Mycrotyloma uniflorum* (with *Solanum tuberosum* as an outgroup) using OrthoFinder^54^ (v2.5.4). This analysis placed 26,038 (84.4%) of the 30,840 AYB proteins into orthogroups. Clustering using Venn Diagrams^55^ revealed 1,296 AYB proteins (4.2%) segregated in 384 species-specific orthogroups (Fig. 7).

**Fig. 7:**
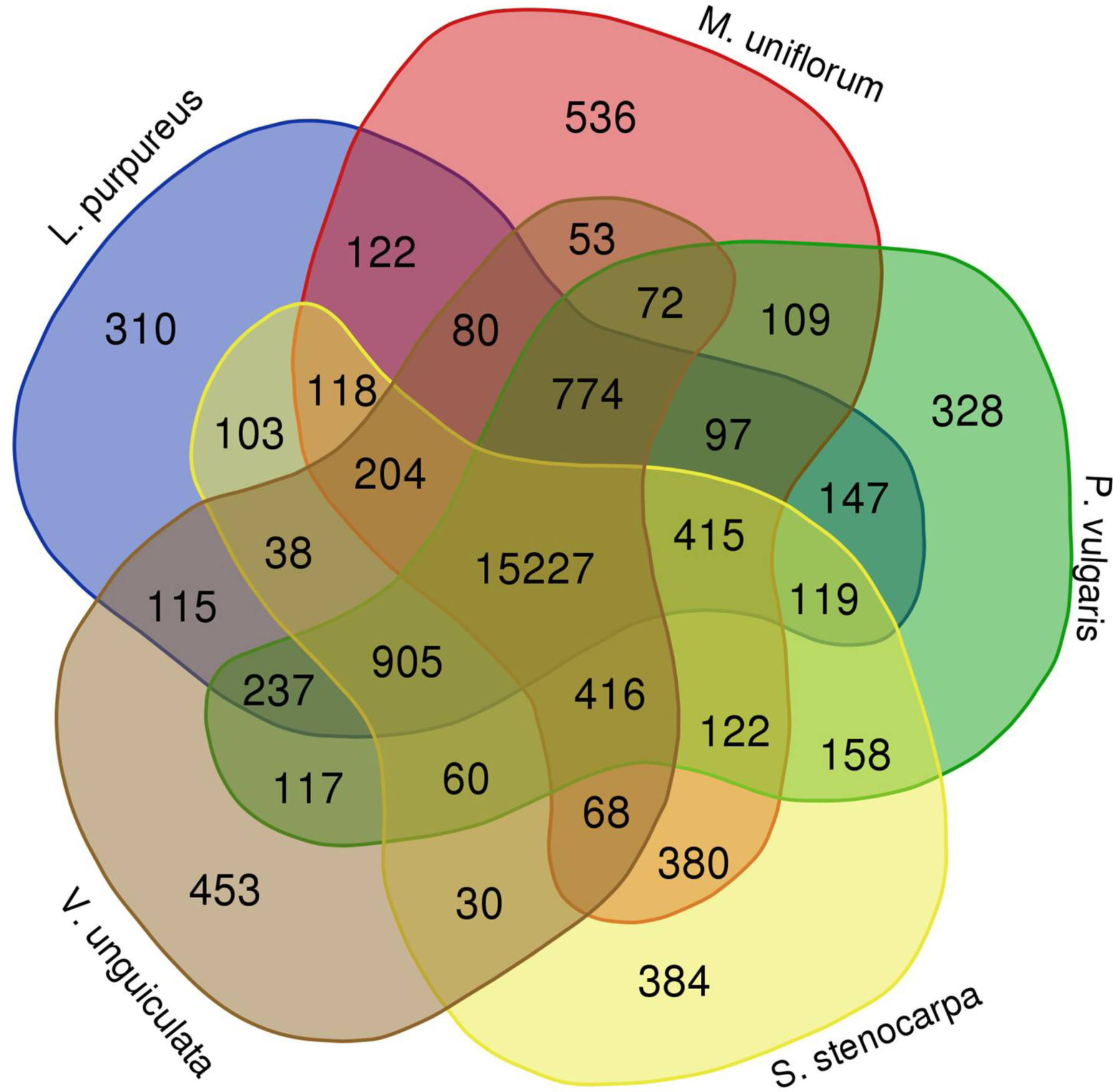
Gene families in *AYB.* Venn diagram of the number of gene families common among and unique to AYB (S. stenocarpa), *Lablab purpureus*, *Phaseolus vulgaris*, *Vigna unguiculata* and *Macrotyloma uniflorum*.

## Technical Validation

### Genome and annotation completeness

We mapped the ONT long-reads data back to the genome assembly and analysed the alignment with Qualimap^56^ (v.2.2.2). The alignment mapping rate was 97.8% (Table 1) with an average read coverage of 39x. We also evaluated the completeness of the genome assembly and annotation using BUSCO^21^ (v5.2.2). A highly conserved set of single-copy orthologs from embryophta_odb10 and fabales lineages were used as references. For the genome assembly, we obtained complete matches to 98.0% and 98.5% of the conserved single-copy orthologs in the fabales and embryophyta lineages, respectively (Fig. 8). Similarly, 90.4% and 91.4% of the conserved single copy orthologs showed complete matches to the gene annotation of AYB (Fig. 8). These high percentages suggest a high degree of accuracy and completeness of the genome assembly and gene annotation.

**Fig. 8:**
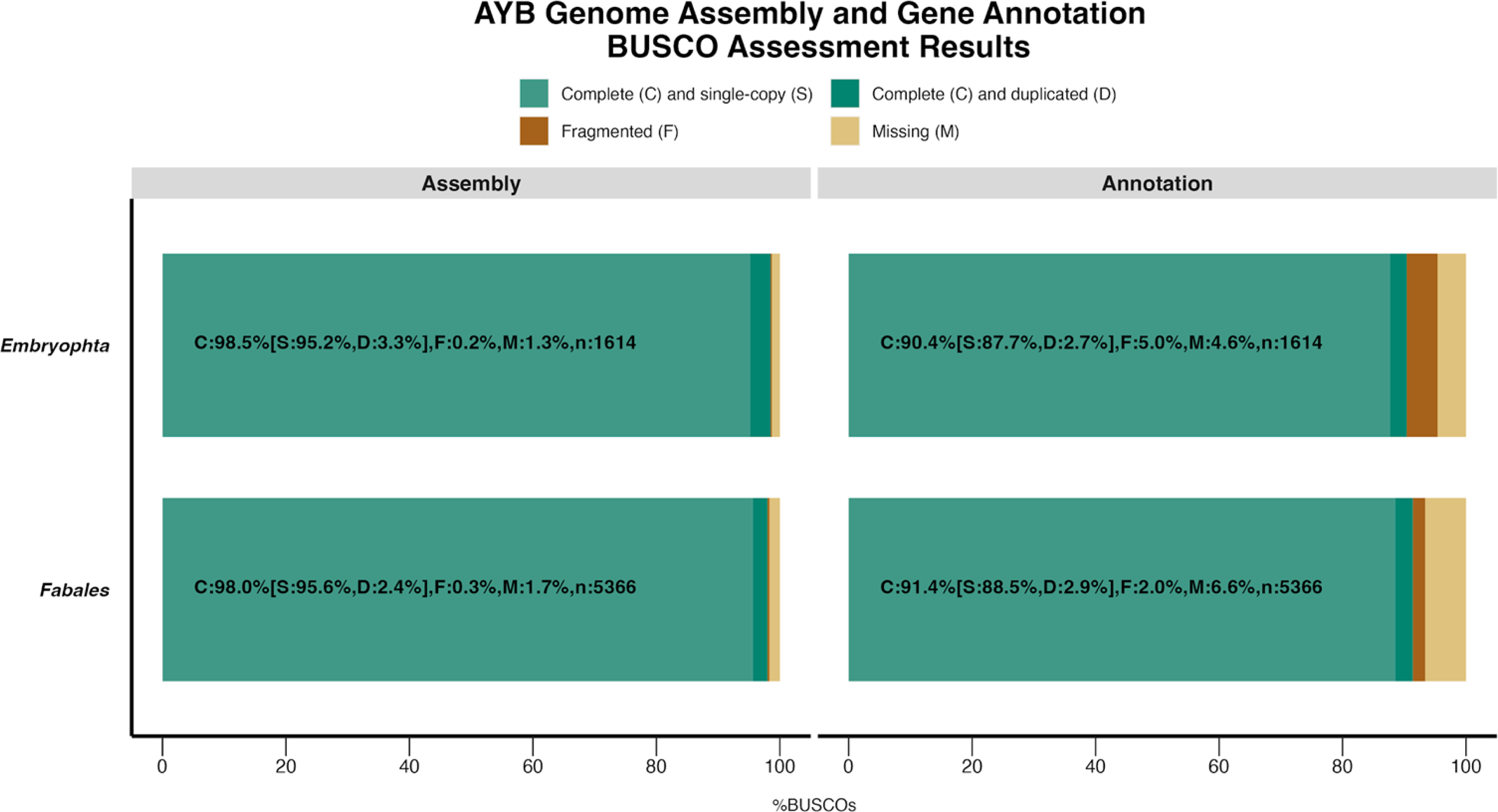
AYB Genome assembly completeness: BUSCO scores of the AYB genome and gene annotation using the embryophyta and fabales reference lineages.

**Fig. 9:**
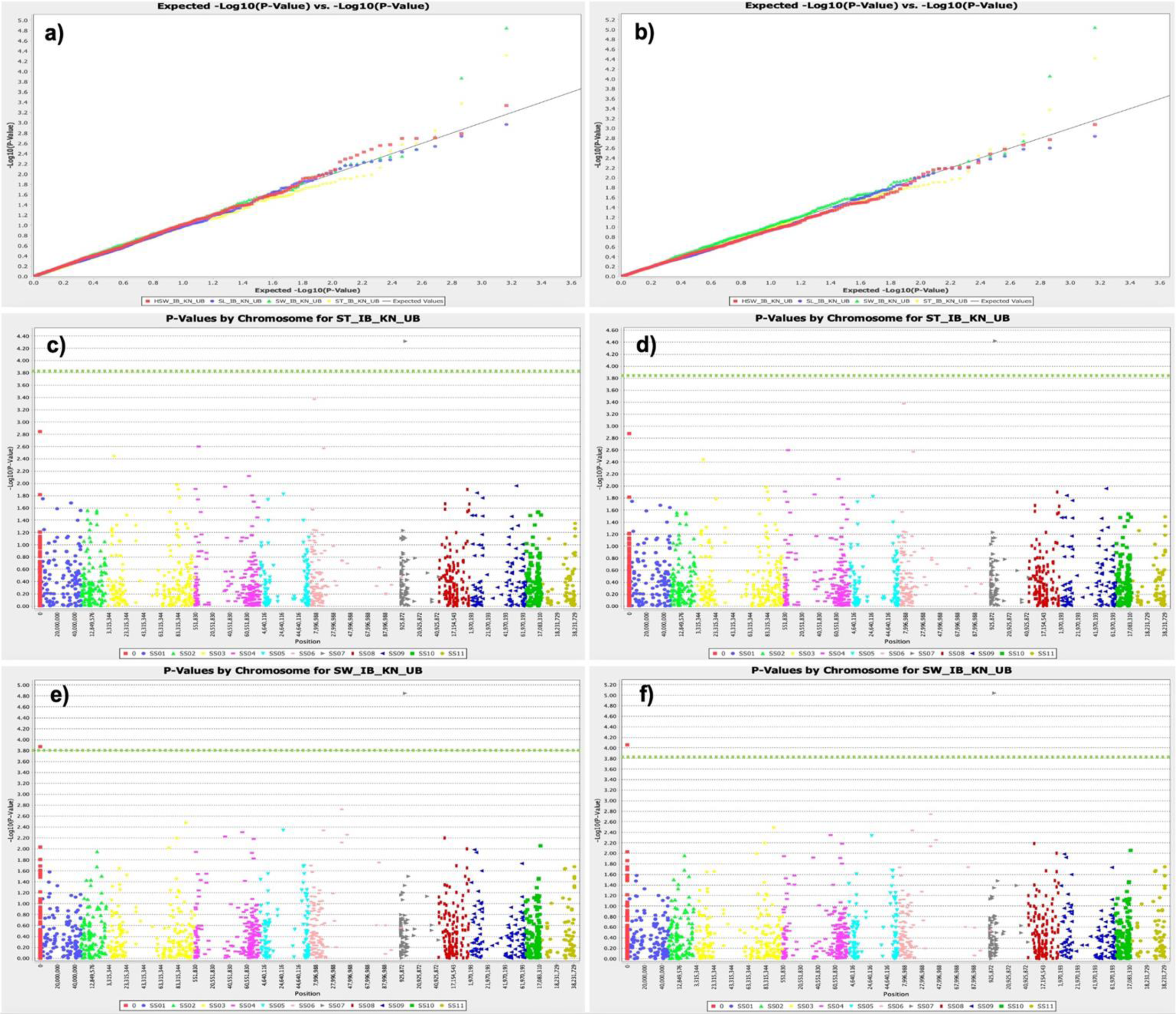
GWAS in AYB. Manhattan and quantile-quantile plot of GWAS analysis for seed thickness (ST) and seed width (SW) in African yam bean collection. (a & b) Q-Q plot for GWAS results for ST and SW traits using GLM and MLM statistical models respectively. (c & d) Manhattan plot for GWAS results of ST trait using GLM and MLM statistical models respectively. (e & f) Manhattan plot for GWAS results of SW trait using GLM and MLM statistical models respectively. The green horizontal dotted lines indicate the significant threshold for associated SNPs in GWAS analysis.

### Marker mapping and association

We also examined the usefulness of the AYB genome for positionally anchoring markers for genetic analyses. Previous efforts to anchor a set of Genotyping-By-Sequencing (GBS) markers generated from a collection of AYB accessions using the common bean genome as reference only mapped 15.48% of the markers to a unique syntenic position, thus limiting the number of markers used for genome-wide association analyses (GWAS)^57^. Using the chromosome-scale assembly of AYB as reference, we could anchor 92% of the total 5,142 DArTseq-SNPs markers to unique positions in the AYB genome. The distribution of the markers across the genome tallies with the gene distribution highlighting the gene-centric nature of the GBS pipeline (Fig. 5). Furthermore, we used a subset of 1,460 quality-filtered SNPs (Call Rate > 0.70, Marker repeatability > 0.95, MAF > 0.05, missing < 0.05) for evaluating how the chromosome-scale genome assembly support GWAS and candidate gene analyses. For this, we used Best Linear Unbiased Estimates (BLUE, combined years and locations) of seed yield traits (hundred seed weight - HSW; seed length -SL; seed width - SW; and seed thickness - ST) from a landrace population of 195 AYB accessions which were phenotyped in 2018 and 2019 under optimal field condition in three different locations of IITA research farms in Nigeria (Ibadan, Kano and Ubiaja). More information about field trait analysis can be obtained from Olomitutu et al (2022)^57^.

The Generalised Linear Model (GLM) and Mixed Linear Model (MLM) were used in TASSEL^58^ (v5.2.87) for identifying marker-trait association for seed yield-traits. Significant SNP association with the traits were determined by adjusting p-value threshold (a = 0.10) for FDR procedure proposed by Benjamini and Hochberg (BH)^59^. Significant marker-trait associations were found for SW and ST traits, two highly correlated traits, on Ss07 and on unanchored contigs. SNP 29420736-57-G/T was associated with both SW and ST traits on Ss07 (4.78 Mbp) of the AYB genome, suggesting a possible pleiotropic effect. The unanchored SNP 29420736-57-G/T was associated with SW traits. The contribution of these associated SNPs to the phenotypic variation ranged between 8.38% to 11.19%.

Candidate genes were searched within 1 Mbp interval around the position of SNP 29420736-57-G/T on Ss07 (± 500 Mb, 4283198 bp to 5283198 bp) in the AYB genome. In total sixteen genes were identified (Supplementary Table 2). Out of 16 candidate genes, nine genes were involving in grain development process in which seven genes were related to seed development role (Spste.TSs11.07G209790.1^60^, Spste.TSs11.07G209800.1^61^, Spste.TSs11.07G209840.1^62^, Spste.TSs11.07G209850.1^62^, Spste.TSs11.07G209860.1^62^, Spste.TSs11.07G209910.1^63^, and Spste.TSs11.07G209920.1^63^), one gene for seed shape (Spste.TSs11.07G209820.1^64,65^), and two genes for seed size (Spste.TSs11.07G209790.1^60^ and Spste.TSs11.07G209920.1^66^) in plants.

Similarly, Olomitutu et al (2022)^57^ reported nine candidate genes in common bean by blasting SNPtag of SNP 29420736-57-G/T in legume information system database^67^. The encoding product of these common bean candidate genes were similarly involved in regulation of seed development^68^, seed/fruit size^69^, seed size^70–73^, grain shape^64,65^, and grain size^74,75^. The mechanism regulating seed traits in AYB needs further exploration. The SNP position and candidate genes information in the AYB genome provided in this study might help to improve AYB yield. These results also indicate that AYB genome will play a central role in precise mapping of SNP markers and genome-wide allele mining for agronomical, biotic, abiotic and nutrition value traits in future AYB crop breeding.

## Data Records

The genome assembly and annotation have been deposited in the following repositories:

1. ENA/NCBI/DDBJ accessions: BioProject PRJEB57813 <u>on ENA</u> and <u>on NCBI</u>, chromosomes have accessions OY731398 to OY731408 and the whole **genome assembly is GCA_963425845.**
2. ENA only: ENA project ERP142818 is already published on INSDC as EDTAPRJEB57813, sample is ERS16321187 (completed) and annotated assembly is ERZ21776326 (completed).

## Code Availability

Open-source software were used for the analyses reported. The software versions and custom parameters used (if different from default) are indicated in the Methods.

## Acknowledgements

This work was supported by the UK Biotechnology and Biological Sciences Research Council through the GCRF Strategic Training Awards for Research Skills (STARS) program grants BB/R020272/1 (ABCF Bioinformatics Community of Practice (BiX COP)) and BB/T017422/1 (Building capacity in third-generation genomics and bioinformatics for agricultural biosciences in Africa). OS was also supported by the Royal Society FLAIR Fellowship (FLR\R1\191850).

## Authors Contribution

OS, JDE, BW, IN, DA, PE, EM, CU, CD, NY and MA designed the experiments. DA, OS, PE, IN, CM, MA, and OO performed the plant sampling. PE, OS, JDE and CM performed genome size estimation, CM, OS, PE, HN, BW, EA, JDE extracted DNA and RNA for sequencing. CM, OS, PE, BW, IN conducted the DNA and RNA sequencing (Illumina and Nanopore), IN, BW, MM, DK, OS, JBE, PE, EM, BBA, HN, BL, HI and EA produced the genome and transcript assemblies. IN and PE performed the repeat annotation. BW, IN, OS, EM and JDE performed the gene annotation. EM, BL, BW, OS, EA did the gene family analyses. OS, PE and HN conducted the synteny analyses. RP, OS, MA, OO and DA did the marker and GWAS analyses. BW, IN, RP, EM, MM, DK, BL, HN, BBA, EA, HI, CU, CD, DA, PE, JDE and OS wrote the manuscript.

