## Supplementary Tables for "Chromosome-scale assembly of the African yam bean genome"

**Supplementary Table 1: AYB genome size estimation based on 3 flow cytometry runs, using *Glycine max* as a standard with assumed genome size range of 1.1-1.15 Gbp.**

|  | soybean_AYB1 | | soybean_AYB2 | | soybean_AYB3 | |  |
| --- | --- | --- | --- | --- | --- | --- | --- |
|  | Mean peak intensity | cv | Mean peak intensity | cv | Mean peak intensity | cv |  |
| **M2 (AYB peak)** | 128781.9 | 1.51 | 125865.9 | 1.9 | 125666.3 | 2.25 |  |
| **M3 (soybean peak)** | 176114.2 | 1.21 | 172886.3 | 1.12 | 171199.8 | 1.15 |  |
| **M4 (double AYB peak)** | 244505.1 | 0.31 | 240296.1 | 1.13 | 238994.3 | 1.09 | **Average ∓ Stdev. (Gbp)** |
| **mean AYB (Gbp)** | 0.822646 | | 0.819031 | | 0.825787 | | 0.822∓0.003 |
| **Lower bound AYB (Gbp)** | 0.804365 | | 0.80083 | | 0.807436 | | 0.804∓0.003 |
| **upper bound AYB (Gbp)** | 0.840927 | | 0.837231 | | 0.844138 | | 0.841∓0.003 |

**Supplementary Table 2: Candidate genes IDs identified near (±500Mb) to associated SNP 29420736-57-G/T in AYB genome.**

| **S.No** | **Chr** | **Start** | **End** | **Gene Name** | **Description of protein matches on NCBI** | **Biological Role** | **References** |
| --- | --- | --- | --- | --- | --- | --- | --- |
| 1 | Ss07 | 428919 | 433677 | Spste.TSs11.07G209780.1 | protein lin-12-like; integrin beta-7 isoform/LOC109816877 | - | - |
| 2 | Ss07 | 435547 | 439708 | Spste.TSs11.07G209790.1 | protein IQ-DOMAIN 14; IQ-DOMAIN 19; IQ-DOMAIN 1; Protein IQ-DOMAIN 31; protein IQ-DOMAIN 34 | Seed development and seed size | Guo et al (2021) *Front. Plant Sci.* **11,** doi: 10.3389/fpls.2020.614851 |
| 3 | Ss07 | 448321 | 448869 | Spste.TSs11.07G209800.1 | Zinc finger C2H2-type; putative transcription factor C2H2 family | Seed development | Puentes-Romero et al (2022) *Plants* **11(15)**, doi: 10.3390/plants11151974 |
| 4 | Ss07 | 453361 | 462062 | Spste.TSs11.07G209810.1 | COP1-interacting protein 7-like | light mediated developmental processes | Xu et al (2015) *PLOS Genetics* **11(12)**, https://doi.org/10.1371/journal.pgen.1005747 |
| 5 | Ss07 | 463157 | 465099 | Spste.TSs11.07G209820.1 | putative protein phosphatase 2C-like protein 44; protein-serine/threonine phosphatase | Seed shape | Hu et al (2012) *J. Integr. Plant Biol*. 54, doi: 10.1111/jipb.12008 ; Wang ET AL (2019) *Front. Plant Sci.* **10**, https://doi.org/10.3389/fpls.2019.00469 |
| 6 | Ss07 | 467485 | 467667 | Spste.TSs11.07G209830.1 | - | - | - |
| 7 | Ss07 | 488752 | 490417 | Spste.TSs11.07G209840.1 | Albumin-2 protein | Seed development | Vigeolas et al (2008) *Plant Physiol.***146(1)**, doi: 10.1104/pp.107.111369 |
| 8 | Ss07 | 491974 | 492660 | Spste.TSs11.07G209850.1 | Albumin-2-like protein | Seed development | Vigeolas et al (2008) *Plant Physiol.***146(1)**, doi: 10.1104/pp.107.111369 |
| 9 | Ss07 | 496092 | 496778 | Spste.TSs11.07G209860.1 | 2S Albumin protein | Seed development | Vigeolas et al (2008) *Plant Physiol.***146(1)**, doi: 10.1104/pp.107.111369 |
| 10 | Ss07 | 504250 | 504321 | Spste.TSs11.07G209870.1 | - | - | - |
| 11 | Ss07 | 509764 | 510915 | Spste.TSs11.07G209880.1 | Glycolipid transfer protein 1 | Lipid transfer | West et al (2008) FEBS J. **275(13),** doi: 10.1111/j.1742-4658.2008.06498.x |
| 12 | Ss07 | 513219 | 514922 | Spste.TSs11.07G209890.1 | glycolipid transfer protein 1 | Lipid transfer | West et al (2008) FEBS J. **275(13),** doi: 10.1111/j.1742-4658.2008.06498.x |
| 13 | Ss07 | 515654 | 518484 | Spste.TSs11.07G209900.1 | - | - | - |
| 14 | Ss07 | 519143 | 520300 | Spste.TSs11.07G209910.1 | KIN14B-interacting protein At4g14310 | Seed development | Tian et al (2021) *BMC Plant Biol.* **210,** https://doi.org/10.1186/s12870-021-02988-6 |
| 15 | Ss07 | 521480 | 523343 | Spste.TSs11.07G209920.1 | KIN14B-interacting protein At4g14310; WD40 repeat-like superfamily protein | Seed development and seed size | Tian et al (2021) *BMC Plant Biol.* **210,** https://doi.org/10.1186/s12870-021-02988-6 ; Siou-Luan and Shin-Lon (2018) *Am. J. Agric*. **6**, doi: 10.11648/j.ajaf.20180602.11 |
| 16 | Ss07 | 524694 | 531580 | Spste.TSs11.07G209930.1 | phosphate transporter PHO1 | Phosphorylation activity | Nussaume et al (2011) *Front. Plant Sci.* **2,** doi: 10.3389/fpls.2011.00083 |
